# Broadly neutralizing antibodies against emerging delta-coronaviruses

**DOI:** 10.1101/2024.03.27.586411

**Authors:** Megi Rexhepaj, Young-Jun Park, Lisa Perruzza, Daniel Asarnow, Mathew Mccallum, Katja Culap, Christian Saliba, Giada Leoni, Alessio Balmelli, Courtney N. Yoshiyama, Miles S. Dickinson, Joel Quispe, Jack Taylor Brown, M. Alejandra Tortorici, Kaitlin R. Sprouse, Ashley L. Taylor, Tyler N Starr, Davide Corti, Fabio Benigni, David Veesler

## Abstract

Porcine deltacoronavirus (PDCoV) spillovers were recently detected in children with acute undifferentiated febrile illness, underscoring recurrent zoonoses of divergent coronaviruses. To date, no vaccines or specific therapeutics are approved for use in humans against PDCoV. To prepare for possible future PDCoV epidemics, we isolated human spike (S)-directed monoclonal antibodies from transgenic mice and found that two of them, designated PD33 and PD41, broadly neutralized a panel of PDCoV variants. Cryo-electron microscopy structures of PD33 and PD41 in complex with the PDCoV receptor-binding domain and S ectodomain trimer provide a blueprint of the epitopes recognized by these mAbs, rationalizing their broad inhibitory activity. We show that both mAbs inhibit PDCoV by competitively interfering with host APN binding to the PDCoV receptor-binding loops, explaining the mechanism of viral neutralization. PD33 and PD41 are candidates for clinical advancement, which could be stockpiled to prepare for possible future PDCoV outbreaks.

---

Coronaviruses (CoVs) are positive-strand enveloped RNA viruses that cause mild to deadly respiratory and enteric disease in avian and mammalian species. *Alpha*-CoVs, 229E^1^ and NL63^2^, along with *Beta-CoVs,* HKU1^3^ and OC43^4^, are endemic in humans and mainly associated with mild respiratory infections (common colds), although some cases lead to severe illness, particularly for children, the elderly and immunocompromised individuals. All three highly pathogenic human CoVs belong to the *Beta*-CoV genus and led to two epidemics, mediated by SARS-CoV-1^5–8^ and MERS-CoV^9^, and the COVID-19 pandemic caused by SARS-CoV-2^10,11^.

CCoV-HuPn-2018 and the closely-related HuCCoV_Z19Haiti, which are canine-feline recombinant alphacoronaviruses, were recently identified in patients with pneumonia or acute respiratory symptoms^12–14^. Porcine delta-coronavirus (PDCoV) was shown to be able to use human APN for cell entry and was recently associated with multiple zoonotic transmission events in children with acute undifferentiated febrile illness^15,16^. These spillovers underscore that more coronaviruses than previously appreciated currently circulate in humans and the zoonotic threats posed by coronaviruses from distinct genera.

The coronavirus spike (S) glycoprotein forms homotrimers anchored in the viral membrane to mediate attachment to host receptors and membrane fusion for initiating infection^17–20^. S plays a key role in modulating host and tissue tropism, zoonotic transmission and pathogenesis^21–23^. Given that S is a main target of antibodies and that neutralizing antibody titers are a correlate of protection against coronaviruses, development of vaccines and therapeutics focuses intensively on this glycoprotein target^24–28^.

Here, we report the isolation of human monoclonal antibodies (mAbs) targeting the PDCoV S receptor-binding domain (RBD) from transgenic mice and show that two of them (PD33 and PD41) broadly neutralize a panel of divergent PDCoV isolates harboring S mutations. Cryo-electron microscopy (cryoEM) analysis of the P33 Fab fragment bound to the PDCoV RBD and of the PD41 Fab bound to a variant PDCoV S trimer provide a blueprint of the interactions mediating recognition and broad neutralization. We demonstrate that both mAbs inhibit APN receptor binding to PDCoV S competitively, explaining their mechanism of broad inhibition. PD33 and PD41 are promising candidates for future clinical advancement individually or as a cocktail for mAb therapy.

## Discovery of broadly neutralizing PDCoV monoclonal antibodies (mAbs)

To identify neutralizing antibodies with therapeutic potential, we immunized Alloy ATX transgenic mice harboring the human immunoglobulin loci encoding for the heavy chain and either the lambda (ATX-GL) or the kappa (ATX-GK) light chain. Two ATX-GL and two ATX-GK mice were immunized alternatively with the PDCoV Illinois strain (IL121_2014) RBD and prefusion-stabilized 2P PDCoV_IL121_2014_ S for a total of 4 doses (**Figure 1A**). Mice were sacrificed 8 days after last boost and peripheral blood, spleen and lymph nodes (LN) were collected and cells freshly isolated. PDCoV_IL121_2014_ RBD-specific memory B cells were selected from immunized mice via fluorescence-assisted cell sorting and VH/VL sequences subsequently retrieved by RT-PCR.

**Figure 1:**
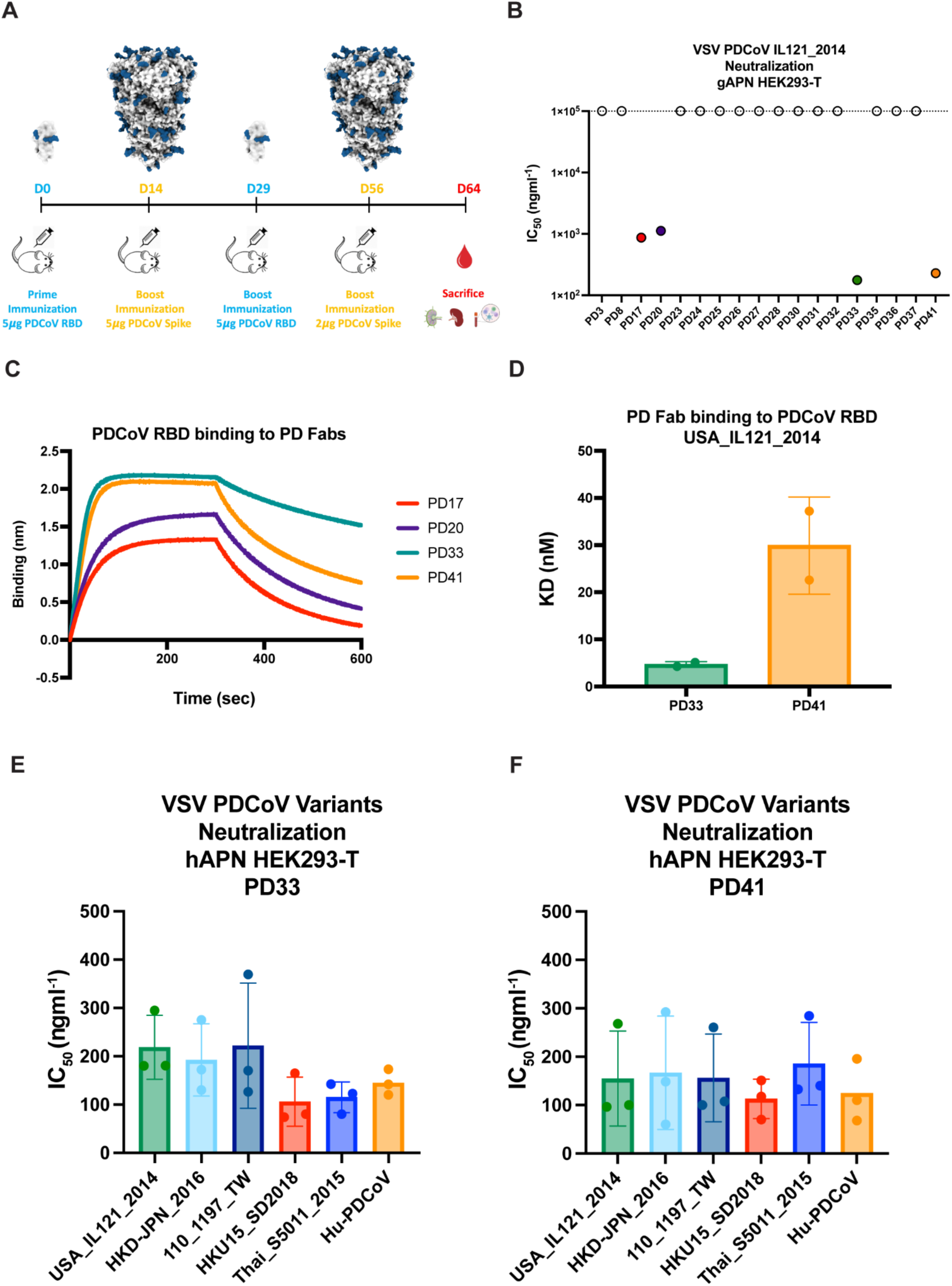
Discovery and characterization of broadly neutralizing PDCoV monoclonal antibodies (mAbs). **A,** Immunization study of ATX mice with PDCoV S and RBD. **B,** Screening of isolated mAbs for neutralization of VSV pseudotyped with PDCoV S using HEK293T cells transfected with the galline APN (gAPN) receptor. Four neutralizing antibodies PD17, PD20, PD33 and PD41 were selected for further characterization. **C,** Biolayer interferometry (BLI) analysis of PDCoV RBD_USA_IL121_2014_ binding to neutralizing antibodies PD17, PD20, PD33, and PD41 Fab fragments. **D,** Binding affinities of the PD33 and the PD41 Fab to PDCoV RBD_USA_IL121_2014_ (dose-response curves are shown in Figure S2). **E-F,** PD33 (E) and PD41 (F) mAb-mediated neutralization of VSV pseudotyped with PDCoV S variants using HEK293T target cells transfected with human APN (hAPN). The bars correspond to geometric mean titers and each data point is one out of 3 biological replicates, each one using a different batch of pseudovirus.

Biolayer interferometry (BLI) and ELISA screening led to the identification of eighteen S-directed mAbs, nine of which binding to the RBD, and all of them were screened for neutralization of a vesicular stomatitis virus (VSV) pseudotyped with PDCoV_IL121_2014_ S using HEK293T target cells transiently transfected with galline APN (gAPN) (**Figure S1A-C**). Four mAbs, designated PD17, PD20, PD33 and PD41, had neutralizing activity with half-maximal inhibitory concentrations (IC_50_) below 1 µg/ml with PD33 and PD41 endowed with the greatest potency (**Figure 1B and Figure S1D**). PD33 and PD41 Fab fragments bound with faster on rates and/or slower off rates to the immobilized PDCoV_IL121_2014_ RBD, relative to PD17 and PD20, with respective binding affinities of ∼5 nM and ∼30 nM (**Figure 1C-D, Figure S2 and Table S1**). Based on these data, PD33 and PD41 were selected as lead PDCoV S-directed antibodies for further characterization.

To investigate the breadth of neutralization mediated by these mAbs, we evaluated their ability to neutralize a panel of VSV pseudotyped with PDCoV S-related isolates harboring mutations in the RBD in presence of the PD33 and the PD41 mAbs using HEK293T target cells transiently transfected with human APN (hAPN). Our panel also included a VSV pseudotyped with the S glycoprotein of the PDCoV isolate identified in humans, which harbor the P8A and V550A residue substitutions. We observed that PD33 and PD41 broadly neutralized our panel of pseudoviruses demonstrating their broad neutralizing activity (**Figure 1E, Figure S3 and Table S2**).

### Molecular basis of broad PD33 mAb-mediated neutralization

To understand the mechanism underlying the broadly neutralizing activity of PD33 towards a panel of PDCoV S variants, we characterized a complex between the PDCoV_IL121_2014_ RBD and the PD33 Fab fragment using cryoEM. To expedite the structural determination process, we opted to use an anti-kappa light chain nanobody recognizing the Fab hinge region and rigidifying it while also augmenting the overall molecular mass^29^. Using this strategy, we determined an asymmetric cryoEM structure at 3.0 Å resolution of this ∼80 kDa complex enabling model building and analysis of the contacts mediating Fab binding (**Figure 2, Figure S4 and Table S3**).

**Figure 2:**
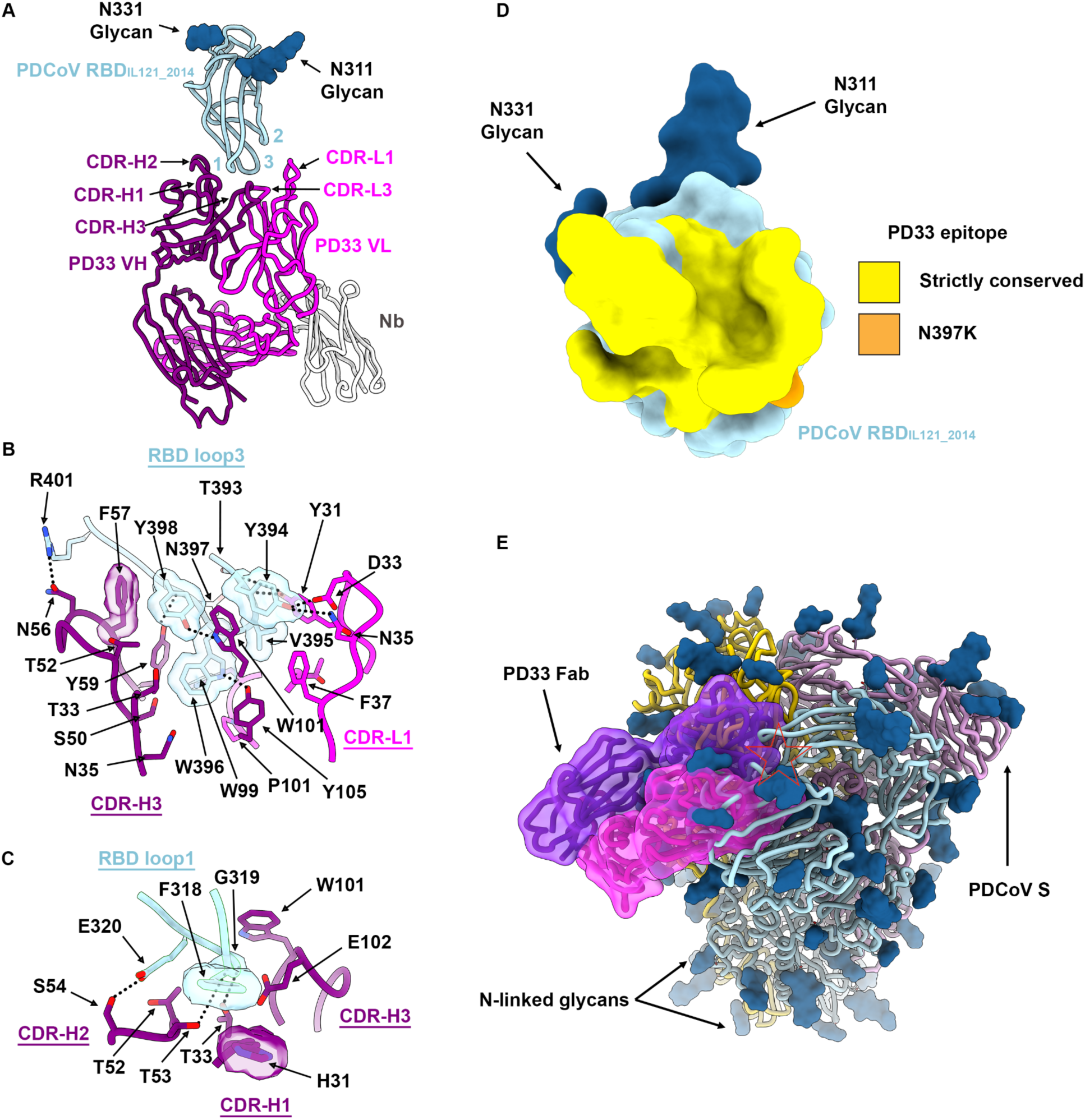
Molecular basis of broad PD33 mAb-mediated neutralization of PDCoV S variants. **A,** Ribbon diagram of the PD33 Fab-bound PDCoV RBD_IL121_2014_ cryoEM structure at 3.0Å resolution. PD33 V_H and_ V_L_ are respectively rendered in purple and pink whereas the PDCoV RBD is colored cyan. The kappa light chain nanobody (Nb) used for assisting structural determination is shown in white. The PDCoV receptor-binding loops are annotated 1-3 and the PD33 complementary determining regions (CDR) for light and heavy chains are annotated CDR-L1, 2, 3 and CHR-H1, 2, 3. N-linked glycans are rendered as blue spheres. **B-C,** Zoomed-in views of the interface between PDCoV RBD_IL121_2014_ loop 3 (B) or loop 1 (C) and the PD33 Fab. Selected hydrogen bonds and salt bridges are shown as black dashed lines. A few side chains are shown in surface representation to highlight shape complementarity. **D,** PD33 epitope conservation across the panel of PDCoV S variants analyzed. Yellow indicates strict residue conservation whereas orange shows the position of the N397K substitution present in S_SD2018/300_. **E,** Superimposition of the PD33-bound PDCoV RBD structure with the PDCoV S ectodomain trimer structure (PDB 6BFU)^35^ showing that PD33 could not bind to a closed S trimer due to masking of the receptor-binding loops and resulting steric clashes (red star).

PD33 binding leads to burial of an average surface of 830 Å^2^ at the interface between the epitope and the paratope through hydrogen bonding and shape complementarity involving the PDCoV RBD loops 1 and 3 and the Fab complementary determining loops (CDRs) H1-3, L1 and L3 (**Figure 2A**). The epitope is dominated by RBD loop 3 which accounts for two thirds of the surface buried at the interface with the PD33 paratope. The PD33 epitope comprises residues 316, 318-322 and 393-401. RBD loop 3 inserts at the interface between PD33 VH and VL through contacts dominated by the W396_IL121_2014_, Y398_IL121_2014_ and R401_IL121_2014_ side chains which are respectively hydrogen bonded to the PD33 VH Y105, W101 and N56 side chains (**Figure 2B**). Furthermore, the Y394_IL121_2014_ phenol is hydrogen bonded to the CDRL1 D33 and N35 side chains whereas the CDRL1 Y31 phenol is hydrogen bonded to the T393_IL121_201_ and Y394_IL121_201_ backbone carbonyls (**Figure 2B**). Extensive van der Waals interactions involving loop 3 Y394_IL121_2014_, V395_IL121_2014_, W396_IL121_2014_ and Y398_IL121_2014_ residues with surrounding CDRH2, CDRH3, CDRL1 and CDRL3 residues further strengthen binding. PD33 CDRH1-3 triangulate RBD loop1 via shape complementarity and hydrogen-bonding of the F318_IL121_2014_ backbone carbonyl and the CDRH2 T53 side chain hydroxyl as well as the E320_IL121_2014_ and the CDRH2 S54 side chains (**Figure 2C**). Strict conservation of epitope residues rationalizes the observed PD33-mediated broad neutralization of PDCoV variants with the S_SD2018/300_ N397K substitution being the sole epitope variability among the strains analyzed (**Figure 2D**). Given that the RBD residue 397 side chain points away from the binding interface and that its backbone is hydrogen-bonded to the PD33 CDRH2 Y59 side chain hydroxyl, the N397K substitution does not affect recognition or neutralization (**Figures 1E and 2B**).

The PD33 epitope comprises residues that are buried and therefore inaccessible in the closed S trimer due to tertiary and quaternary interactions (**Figure 2D**). As a result, PD33 binding requires RBD opening, as it would be incompatible with the closed RBD conformation. This is reminiscent of some SARS-CoV-1, SARS-CoV-2 and MERS-CoV antibodies that conformationally select for open RBDs and trigger S_1_ shedding and refolding to the postfusion conformation via receptor-functional mimicry^30–34^.

### Molecular basis of broad PD41 mAb-mediated neutralization

To obtain a detailed understanding of PD41 mAb-mediated neutralization of a large panel of PDCoV-related isolates, we selected a PDCoV variant isolated from South Dakota in 2018 (S_SD2018_) which harbor three RBD mutations (M354I, I391V, N397K) relative to RBD_IL121_2014_. S_SD2018_ VSV exhibited the most efficient entry into HEK293T cells transiently transfected with human APN among the 6 isolates tested (**Figure S5**). Furthermore, the RBD_SD2018_ bound to human APN with a 5-fold greater affinity than that of RBD_IL121_2014_, due to modulation of off-rates, similar to what has been observed for SARS-CoV-2 variants^36–39^.

We characterized the complex between the PD41 Fab fragment and the S_SD2018_ ectodomain trimer using cryoEM and identified four distinct conformational states of the S glycoprotein by 3D classification : (i) class I with all three RBDs closed without any bound Fabs; (ii) class II with one Fab-bound RBD; (iii) class III with two Fab-bound RBDs, (iv) and class IV with three Fab-bound RBDs (**Figure 3A, Figure S6 and Table S3**). To improve the resolvability of the PD41/RBD interface, we used local refinement of class II to obtain a map focused on this region at 3.0Å resolution (**Figure S6**).

**Figure 3.**
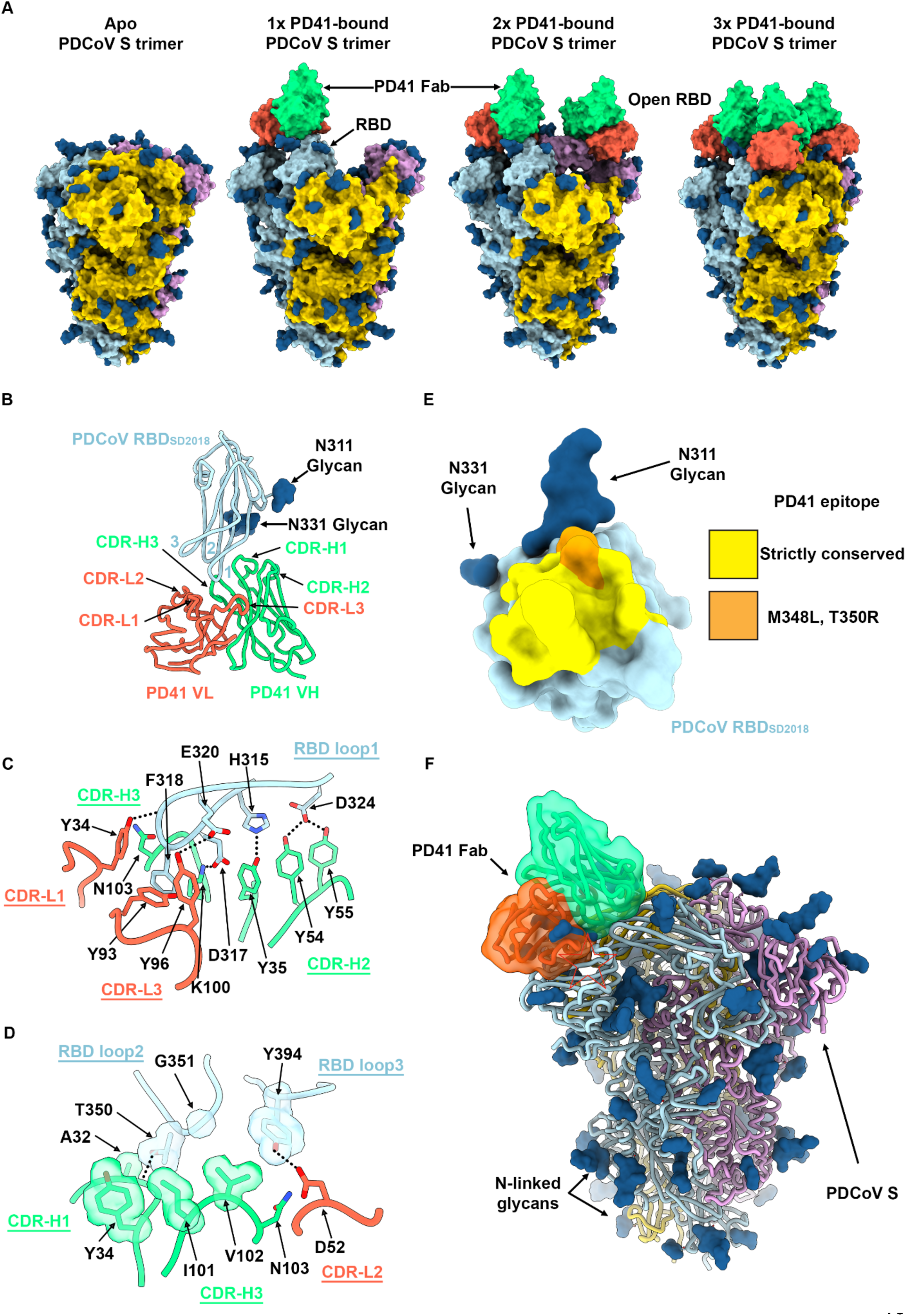
Molecular basis of broad PD41 mAb-mediated neutralization of PDCoV S variants. **A,** Surface renderings of the PDCoV S_SD2018_ trimer (gold, cyan and pink) without and with one, two and three bound PD41 Fabs (green and orange for heavy and light chains, respectively) observed by cryoEM. N-linked glycans are rendered as blue spheres. **B,** Ribbon diagram of the PD41 Fab-bound PDCoV RBD_SD2018_ cryoEM structure at 3.0Å resolution obtained through local refinement of the one PD41-Fab bound S structure. PD41 V_H and_ V_L_ are respectively rendered in green and orange whereas the PDCoV RBD is colored cyan. The PDCoV receptor-binding loops are annotated 1-3 and the PD41 complementary determining regions (CDR) for light and heavy chains are annotated CDR-L1, 2, 3 and CHR-H1, 2, 3. **C-D,** Zoomed-in views of the interface between PDCoV RBD_SD2018_ loop 1 (C) or loops 2 and 3 (D) and the PD41 Fab. Selected hydrogen bonds and salt bridges are shown as black dashed lines. A few side chains are shown in surface representation to highlight shape complementarity. **E,** PD41 epitope conservation across the panel of PDCoV S variants analyzed. Yellow indicates strict residue conservation whereas orange shows the positions of the M348L and T350R substitutions present in S_Thailand/S5011/2015_. **F,** Superimposition of the PD41-bound PDCoV RBD structure with the PDCoV S ectodomain trimer structure (PDB 6BFU)^35^ showing that PD41 could not bind to a closed S trimer without structural changes due to masking of the receptor-binding loops and resulting steric clashes (red star).

PD41 binding leads to burial of 615 Å^2^ of the epitope and of the paratope surfaces through contacts involving all six CDR loops and the three PDCoV RBD loops through a combination of hydrogen bonds, salt bridges and shape complementarity (**Figure 3B)**. CDR H1 and H3 dominate the interface and account for two thirds of the buried surface upon PD41 binding along with most polar interactions. The PD41 epitope comprises residues 313-320, 322, 324, 348, 350-351 and 394. The RBD_SD2018_ inserts loop 1 F318_SD2018_ side chain in a hydrophobic cavity formed at the interface between the PD41 VH and VL domains and its backbone amide group forms hydrogen bonds with the VH V102 backbone carbonyl and with the VH N103 side chain amide (**Figure 3B-C)**. Furthermore, RBD_SD2018_ loop 1 residues H315, D317, E320 and D324 form a hydrogen bond and salt bridge network with the CDRH1 Y35, CDRH2 Y54 and Y55, CDRH3 K100 and CDRL3 Y96 and Y99 side chains. Relative to loop 1, RBD loops 2 and 3 make more modest contributions to the epitope through contacts mediated by T350_SD2018_ and G351_SD2018_ as well as Y394_SD2018_, respectively **(Figure 3D)**. Strict conservation of epitope residues rationalizes the observed PD41-mediated broad neutralization with the S_Thailand/S5011/2015_ M348K and T350R substitutions being the sole epitope variability among the variants analyzed (**Figure 3E**). Although the M348 side chain points away from the interface with PD41, the T350 side chain is part of the interface and likely accommodated due to retained neutralization of this variant (**Figure 3D-E**).

The PD41 epitope comprises residues that are masked through tertiary and quaternary interactions in the context of the prefusion closed S trimer^35^ and would preclude Fab binding (**Figure 3F**). As a result, PD41 binding requires structural rearrangements of the S trimer, such as opening of neighboring RBDs (as observed in our cryoEM structures with one and two bound PD41 Fabs) along with relative repositioning of the RBD and the NTD (as is the case in all our PD41 Fab-bound structures) (**Figure 3A**).

### The PD33 and PD41 mAbs competitively inhibit receptor binding

Given that PD33 and PD41 recognize epitopes mapping to the PDCoV S receptor binding loops, we hypothesized that the two mAbs inhibit APN engagement to the RBD competitively. Superimposition of the human APN-bound PDCoV RBD structure to our cryoEM structures further supported this hypothesis by revealing that APN binding would not be compatible with binding of either of these two mAbs due to major steric incompatibilities despite the different angles of approach of each Fab which recognize overlapping but distinct epitopes (**Figure 4-AB and Figure S7)**.

**Figure 4:**
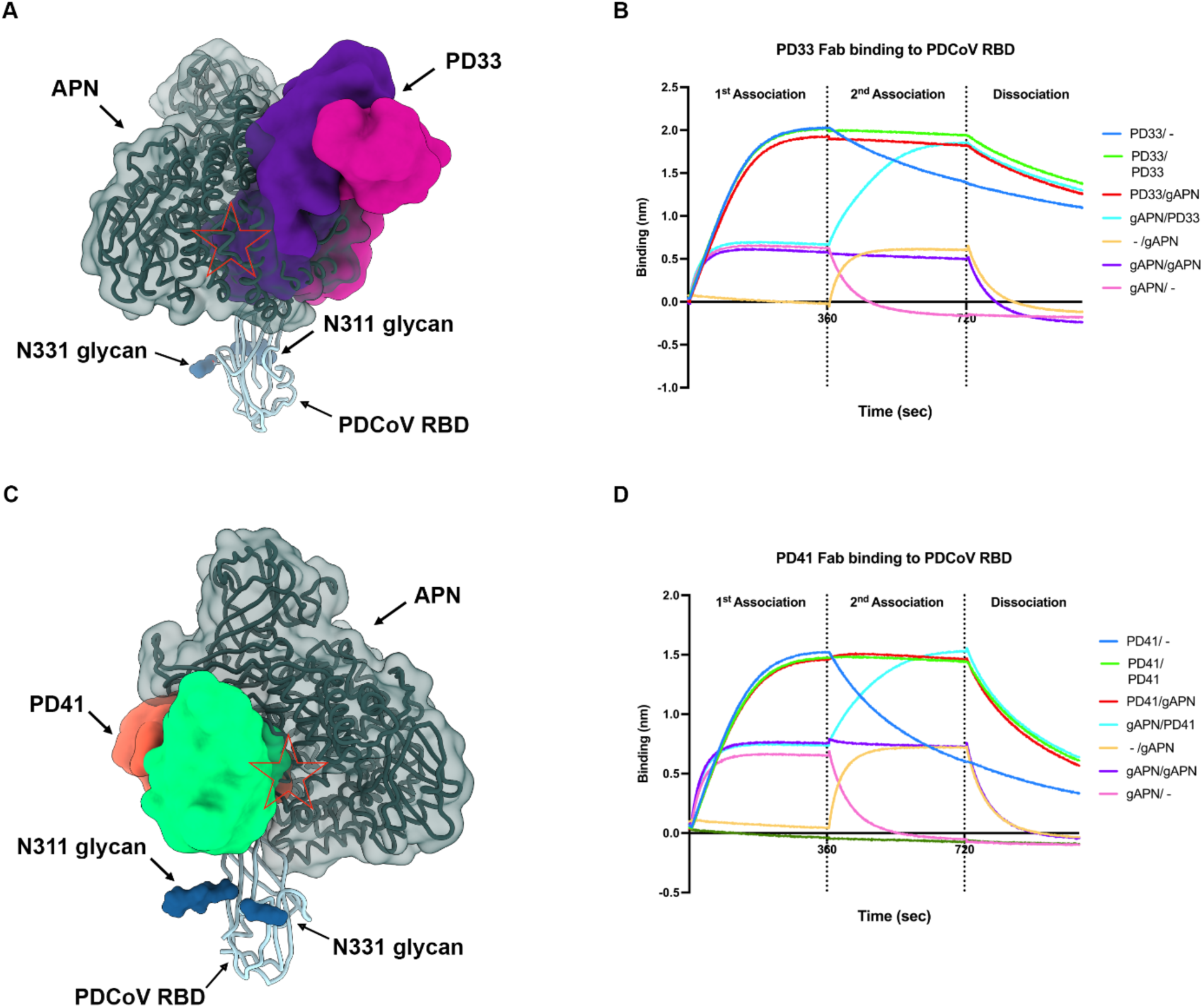
The PD33 and PD41 mAbs competitively inhibit receptor binding to the PDCoV RBD. **A,** PD33 (purple and magenta surfaces for the heavy and light chains, respectively) and human APN (hAPN, dark green) recognize overlapping sites on the PDCoV RBD (cyan ribbon). N-linked glycans are rendered as blue surfaces. The red star indicates steric clashes. **B,** BLI analysis of Fab PD33 binding to the PDCoV RBD immobilized on biosensors in the presence and absence of galline APN (gAPN). **C,** PD41 (purple and magenta surfaces for the heavy and light chains, respectively) and hAPN (dark green) recognize overlapping sites on the PDCoV RBD (cyan ribbon). N-linked glycans are rendered as blue surfaces. The red star indicates steric clashes. **D,** BLI analysis of Fab PD41 binding to PDCoV RBD immobilized on biosensors in the presence and absence of gAPN.

To evaluate the impact of each mAb on APN recognition, we performed a competition experiment to assess APN binding to the PDCoV RBD immobilized at the surface of biolayer interferometry biosensors with or without prior incubation with the PD33 Fab or the PD41 Fab. We found that PD33 or PD41 binding blocked subsequent APN binding to the RBD, thereby confirming the competitive mechanism of inhibition. The reverse experiment, however, revealed that pre-incubation of the immobilized PDCoV RBD with APN did not prevent subsequent Fab binding, suggesting that PD33 and PD41 can displace bound APN, likely due to their higher binding affinity.

## Discussion

The recently described zoonotic introductions and detection of PDCoV in Haitian children^16^ underscores that more coronaviruses spillover to humans than previously appreciated. PDCoV belongs to the most recently described delta-coronavirus genus, which was not previously known to comprise human pathogens, for which correlates of protection are poorly understood. Strikingly, only 4 out of 9 RBD-directed mAbs screened in this study had detectable neutralizing activity against the vaccine-matched pseudovirus and the two most potent mAb inhibited receptor-binding competitively. The PDCoV RBD is thus a prime target of humoral immunity which is reminiscent of SARS-CoV-2 RBD-directed antibodies that account for most of the neutralizing activity against vaccine-matched and mismatched viruses upon infection or vaccination^26,40–42^. Future studies will elucidate the role of NTD-directed and fusion machinery-directed antibodies against PDCoV. The broad neutralizing activity of PD33 and PD41 against a panel of PDCoV isolates suggest that these antibodies would provide some resilience to viral evolution and could be stockpiled for pandemic preparedness and deployed as individual mAb therapies or mAb cocktail.

Although cryoEM structures of the SARS-CoV-1, SARS-CoV-2 and MERS-CoV S trimers revealed the conformational landscape of the RBDs^20,30,43–46^, which can adopt open and closed conformations, other S trimers have largely been found in the closed state^13,18,35,47–52^. Recent work, however, showed that α2,8-linked 9-O-acetylated disialosides binding to the HKU1 S NTD induces RBD opening and subsequent binding to the TMPRSS2 receptor^53,54^. These data suggest that other coronavirus S trimers, such as PDCoV S, could possibly open in response to engagement of an attachment factor, thereby allowing binding to an entry receptor and viral entry. This strategy would promote masking of key sites of vulnerabilities (e.g. the receptor-binding motif) from the immune system while ensuring proper spatial and temporal coordination of the conformational changes required to engage the entry receptor and initiate infection^52^. Future identification of the molecular determinants of PDCoV S opening will reveal the cascade of events leading to productive infection, inform tissue and species tropism and guide vaccine design against this emerging pathogen.

## Supplementary Figures

**Figure S1.**
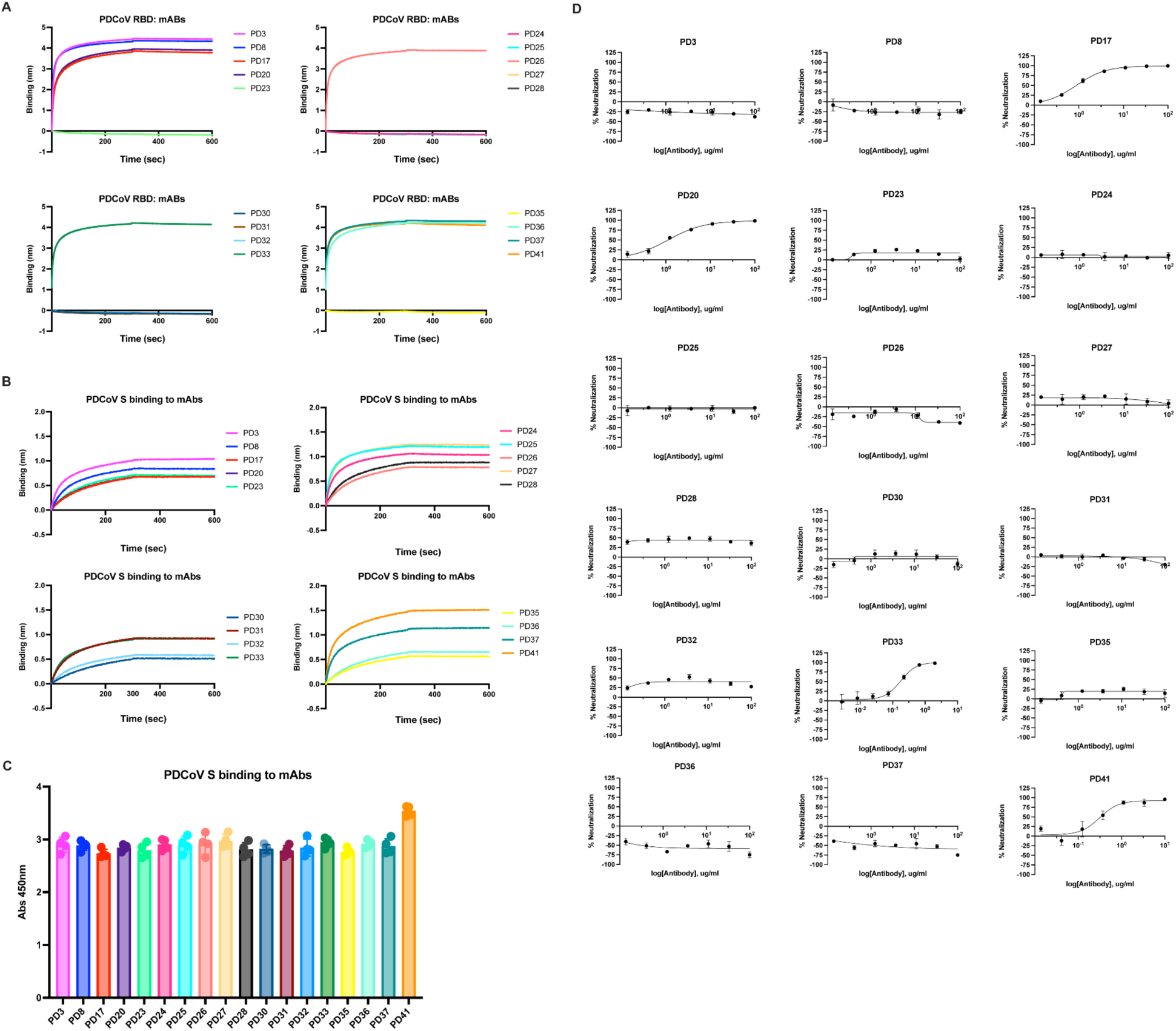
Screening for binding and neutralization of a panel of human mAbs (PD3, PD8, PD17, PD20, PD23, PD24, PD25, PD26, PD27, PD28, PD30, PD31, PD32, PD33, PD35, PD36, PD37, PD41) isolated from transgenic mice. **A,** Screening of isolated mAbs for binding to PDCoV RBD immobilized at the surface of BLI biosensors. **B,** Screening of isolated mAbs immobilized at the surface of BLI biosensors for binding to PDCoV S **C,** Evaluation of binding of the isolated mAbs to PDCoV S measured by ELISA. Each point represents the mean of technical quadruplets. Standard deviations shown as error bars. **D,** Dose-dependent mAb-mediated neutralization of PDCoV S VSV using HEK293T target cells transiently transfected with galline APN (gAPN). Each point represents the mean of technical triplicates. Standard deviations shown as error bars.

**Figure S2:**
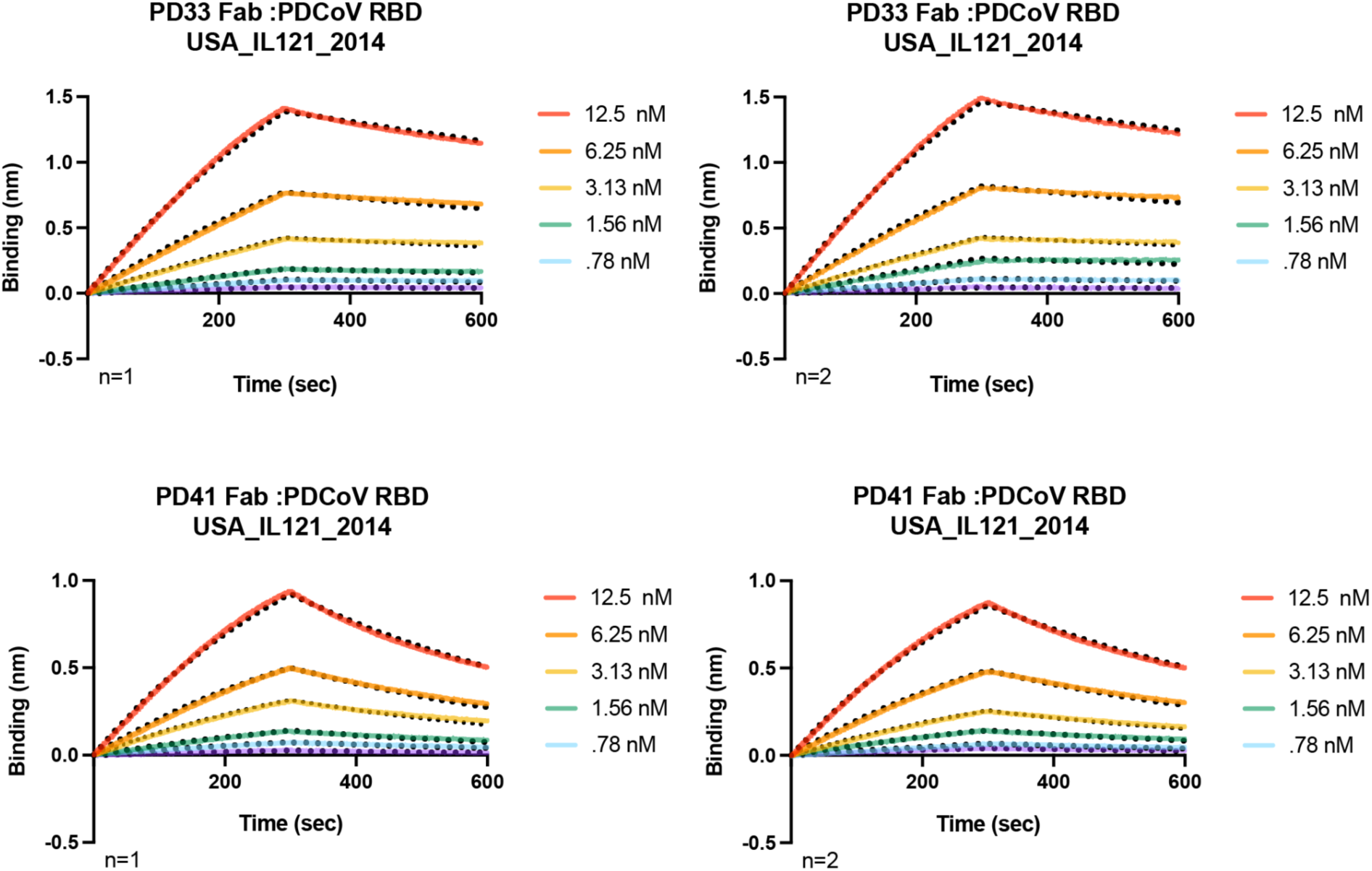
Kinetics of PD33 and PD41 Fabs binding to PDCoV RBD. BLI analysis of PD33 and PD41 Fab fragments binding to immobilized PDCoV_IL121_2014_ RBD immobilized onto Ni-NTA tips. Fits to the data are shown as black dotted lines and were used to determine the binding affinity (K_D_) of the PD33 and PD41 Fab fragments to PDCoV RBD. Two biological replicates are shown for each Fab.

**Figure S3.**
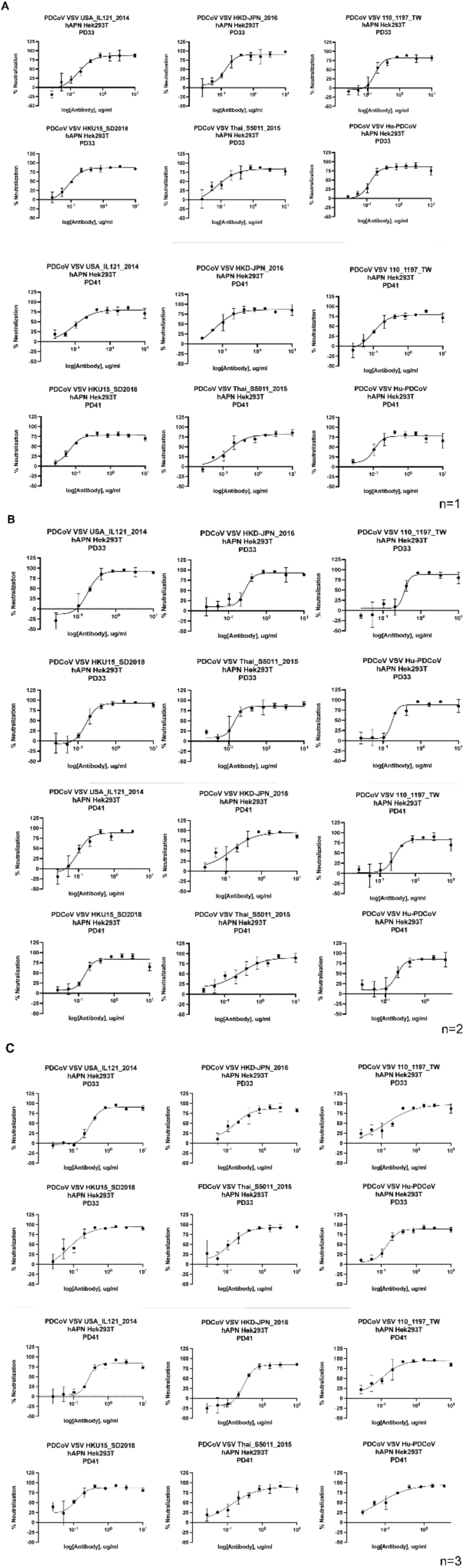
Assessment of PD33 and PD41 neutralization breadth. Dose-dependent mAb-mediated neutralization of PDCoV S VSV variants using HEK293T target cells transiently transfected with human APN. A-C are three independent runs with 3 different batches of pseudovirus.

**Figure S4:**
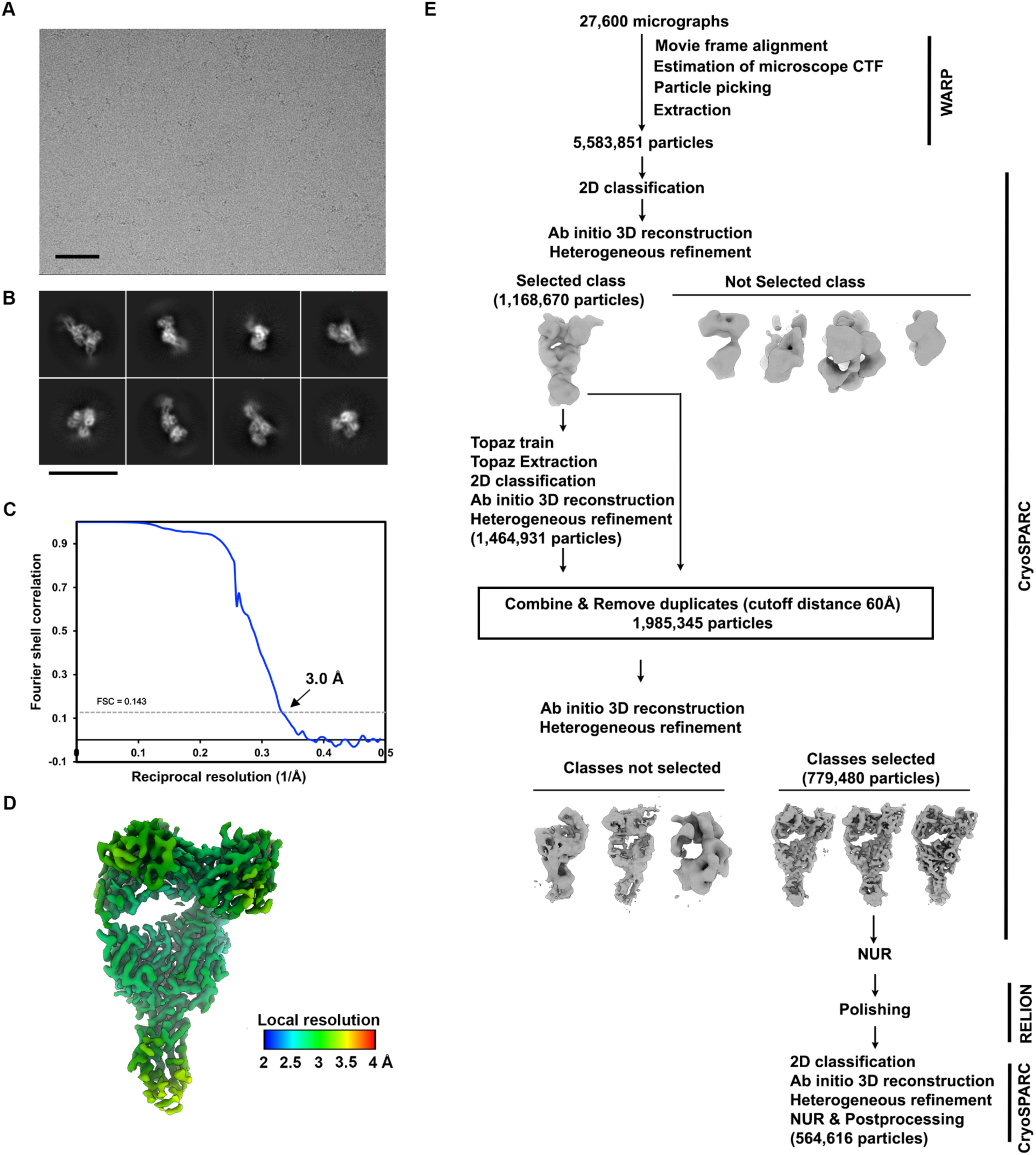
Cryo-EM data collection and refinement of PDCoV RBD_IL2014_ bound to the PD33 Fab fragment. **A,** Representative electron micrograph (3.0 µm defocus, scale bar = 100 nm) **B,** 2D class averages (scale bar = 150 Å). **C,** Gold-standard Fourier shell correlation curve for the cryoEM reconstruction. The 0.143 cutoff is indicated with a gray dashed line. **D,** 3D reconstruction of PDCoV RBD bound to PD33 colored by local resolution calculated using CryoSPARC. **E,** Flow chart of the pipeline for processing. CTF: contrast transfer function; NUR: non-uniform refinement.

**Figure S5.**
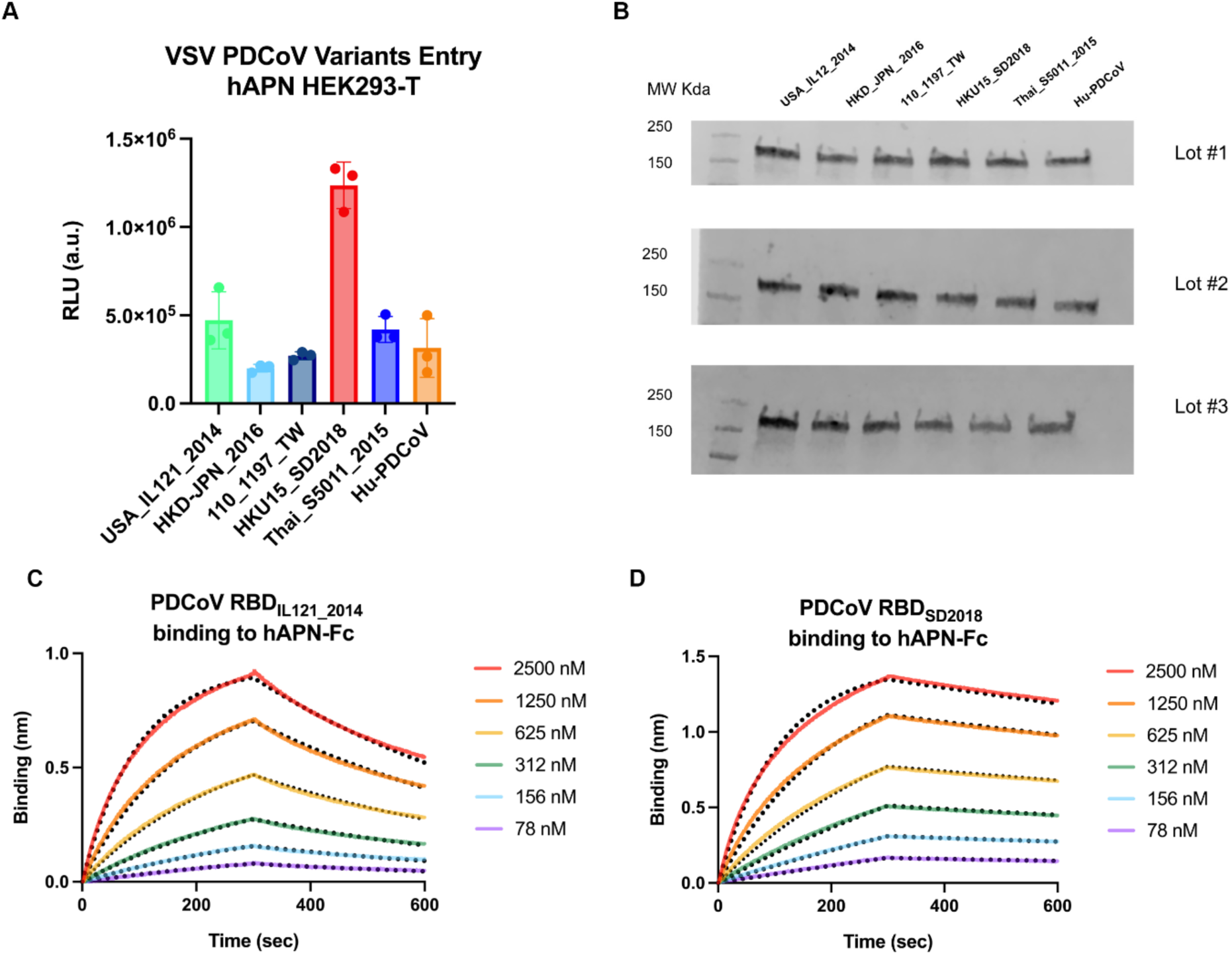
Functional characterization of PDCoV S variants. **A,** Entry of PDCoV S VSV variants into HEK293T target cells transiently transfected with human APN. **B,** Western blot quantification of PDCoV S incorporation in VSV pseudotypes for each of the three biological replicates used in panel A. **C-D,** BLI analysis of hAPN-Fc binding to the PDCoV_IL121_2014_ RBD (left) or to the PDCoV_SD_2018_ (right) immobilized onto Ni-NTA tips. Fits to the data are shown as black dotted lines and were used to determine the apparent binding affinity (K_D,app_) of hAPN-Fc fragments to the two PDCoV RBDs.

**Figure S6:**
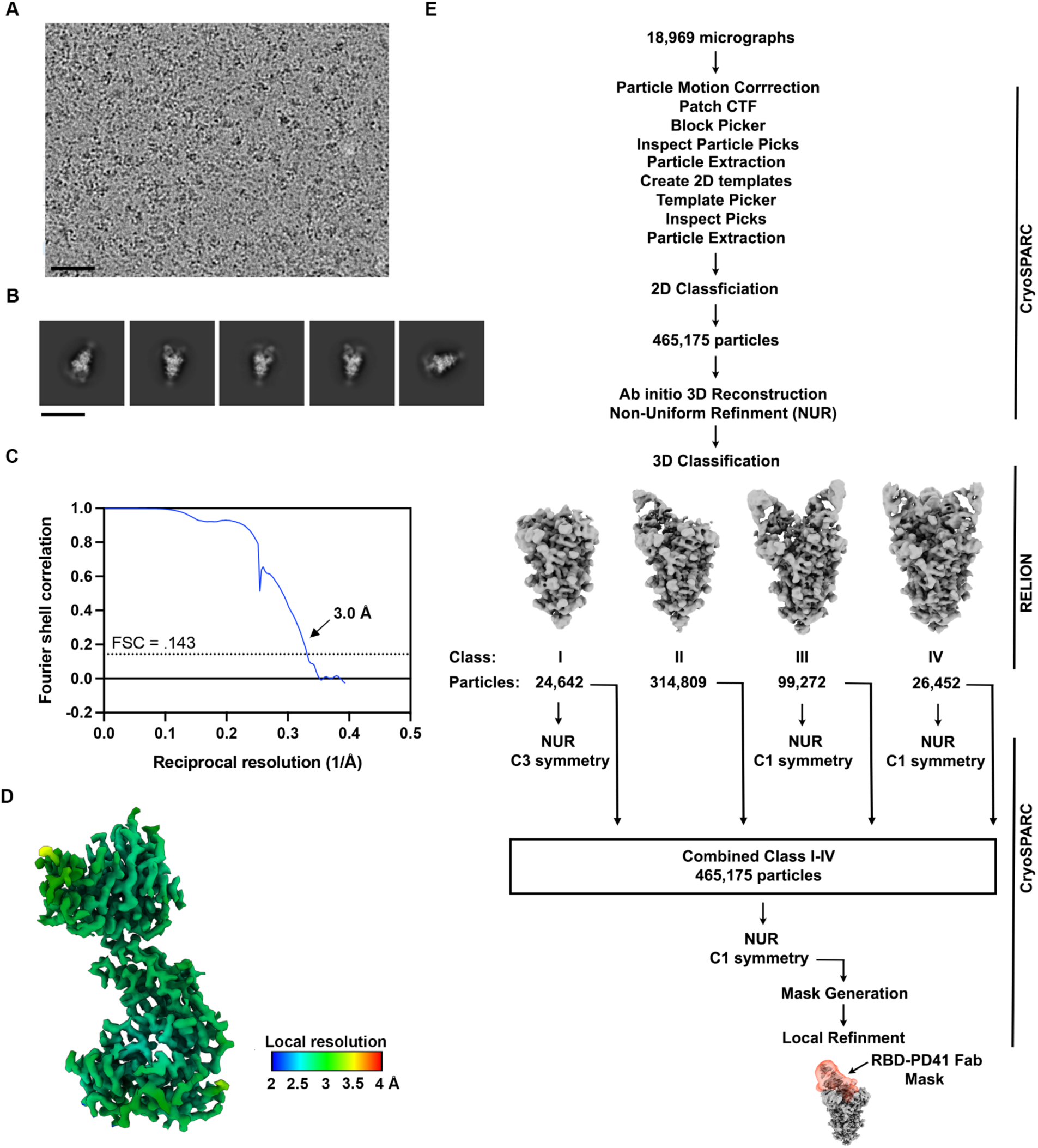
Cryo-EM data collection and refinement of PDCoV RBD_SD2018_ bound to the PD41 Fab fragment. **A,** Representative electron micrograph (2.5 µm defocus, scale bar = 100 nm) **B,** 2D class averages (scale bar = 150 Å). **C,** Gold-standard Fourier shell correlation curve for the cryoEM reconstruction. The 0.143 cutoff is indicated with a black dashed line. **D,** 3D reconstruction of PDCoV RBD bound to PD41 colored by local resolution calculated using CryoSPARC. **E,** Flow chart of the pipeline for processing. CTF: contrast transfer function; NUR: non-uniform refinement.

**Figure S7:**
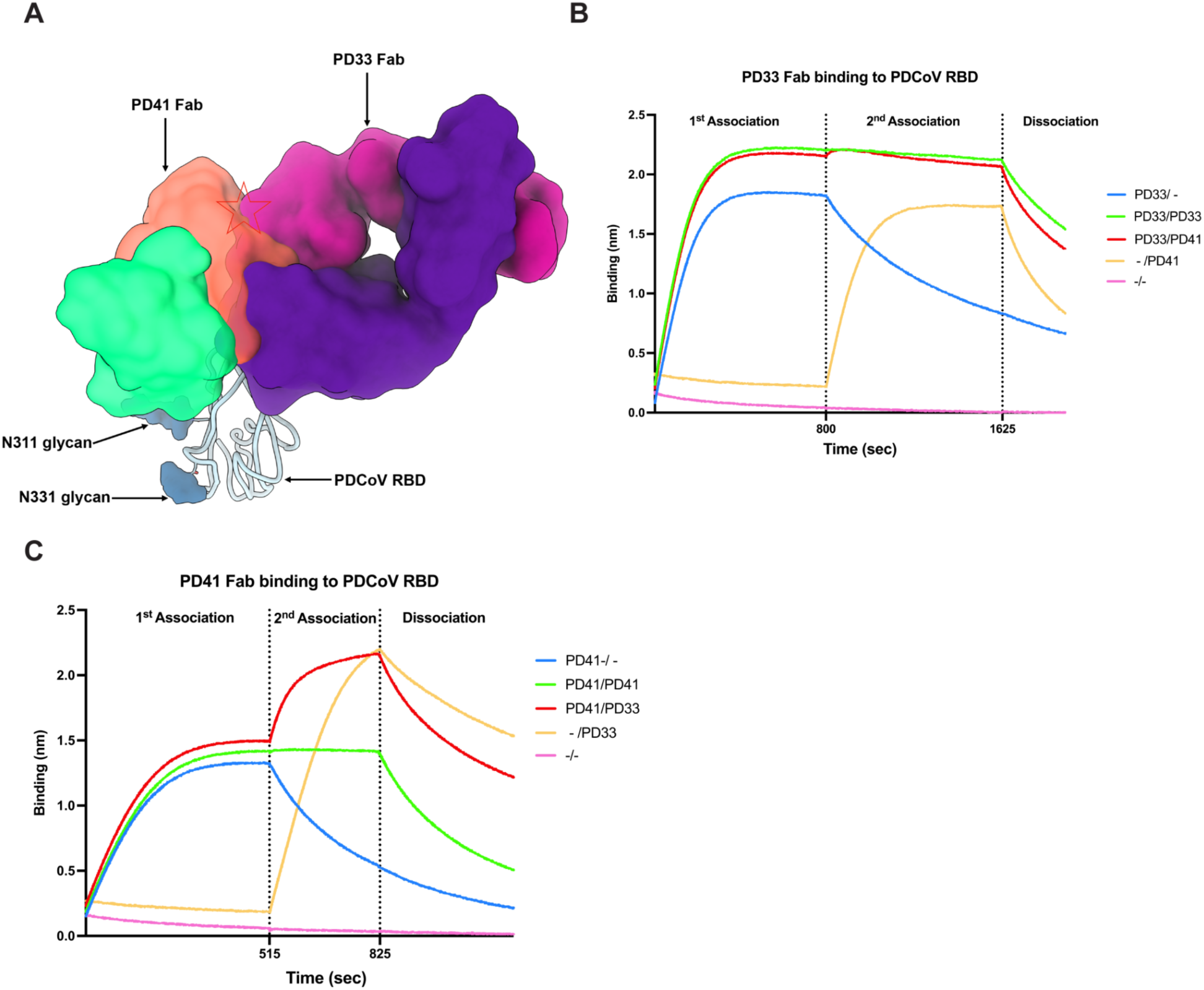
Superimposition of the PD33-bound PDCoV RBD and of the PD41-bound PDCoV RBD_SD2018_ structures showing recognition of distinct but overlapping epitopes. **A,** PD33 (purple and magenta surfaces for the heavy and light chains, respectively) and PD41 (green and orange surfaces for the heavy and light chains, respectively) recognize overlapping epitopes on the PDCoV RBD (cyan ribbon). N-linked glycans are rendered as blue surfaces. The red star indicates steric clashes. **B,** BLI analysis of Fab PD33 binding to the PDCoV RBD immobilized on biosensors in the presence and absence of Fab PD41. **C,** BLI analysis of Fab PD41 binding to the PDCoV RBD immobilized on biosensors in the presence and absence of Fab PD33.

**Table S1.**
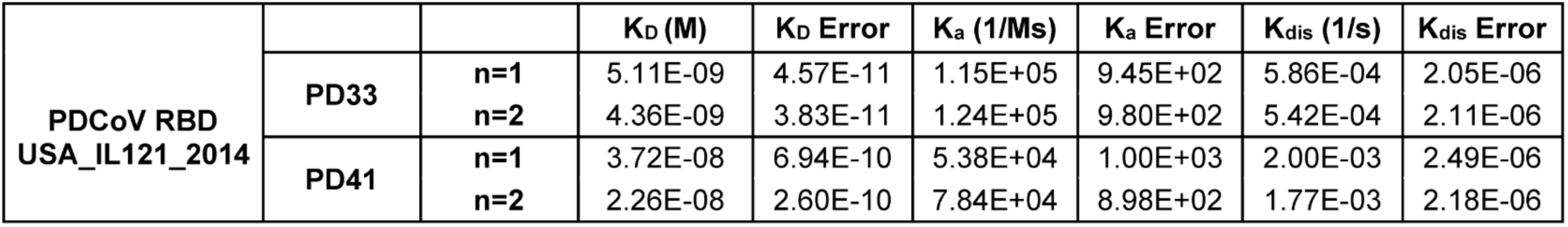
Table of the KDs, association (Ka) and dissociation rate constants (Kdis) for duplicate lots of PDCoV RBD_IL121_2014_ and Fab fragments of PD33 and PD41.

**Table S2.**
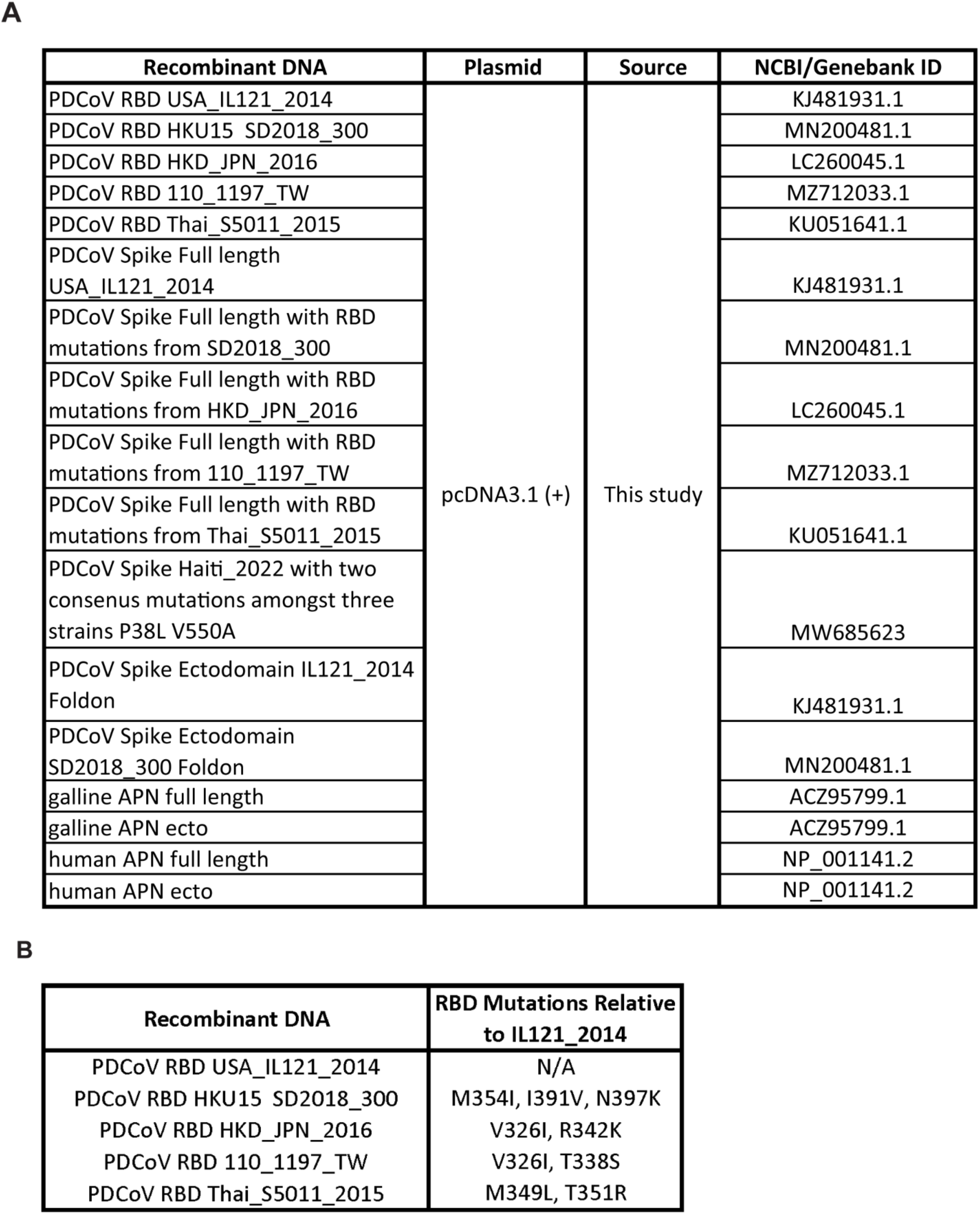
Table of the sequence information of PDCoV S variants as full length, ectodomain and RBD along with hAPN and gAPN sequences as full length and ectodomain constructs. **A,** List of PDCoV variants used in the study and the genebank ID associated for those variants. The table includes information for the APN constructs used in the study and the genebank ID corresponding to those constructs. **B,** List of the PDCoV variants tested in the study and the RBD mutations found in these variants relative to PDCoV_IL121_2014_.

**A**
| Recombinant DNA | Plasmid | Source | NCBI/Genebank ID |
| --- | --- | --- | --- |
| PDCoV RBD USA_IL121_2014 | pcDNA3.1 (+) | This study | KJ481931.1 |
| PDCoV RBD HKU15_SD2018_300 |  |  | MN200481.1 |
| PDCoV RBD HKD_JPN_2016 |  |  | LC260045.1 |
| PDCoV RBD 110_1197_TW |  |  | MZ712033.1 |
| PDCoV RBD Thai_S5011_2015 |  |  | KU051641.1 |
| PDCoV Spike Full length USA_IL121_2014 |  |  | KJ481931.1 |
| PDCoV Spike Full length with RBD mutations from SD2018_300 |  |  | MN200481.1 |
| PDCoV Spike Full length with RBD mutations from HKD_JPN_2016 |  |  | LC260045.1 |
| PDCoV Spike Full length with RBD mutations from 110_1197_TW |  |  | MZ712033.1 |
| PDCoV Spike Full length with RBD mutations from Thai_S5011_2015 |  |  | KU051641.1 |
| PDCoV Spike Haiti_2022 with two consensus mutations amongst three strains P38L V550A |  |  | MW685623 |
| PDCoV Spike Ectodomain IL121_2014 Foldon |  |  | KJ481931.1 |
| PDCoV Spike Ectodomain SD2018_300 Foldon |  |  | MN200481.1 |
| galline APN full length |  |  | ACZ95799.1 |
| galline APN ecto |  |  | ACZ95799.1 |
| human APN full length |  |  | NP_001141.2 |
| human APN ecto |  |  | NP_001141.2 |

**Table S2. Table of the sequence information of PDCoV S variants as full length, ectodomain and RBD along with hAPN and gAPN sequences as full length and ectodomain constructs.**
| Recombinant DNA | RBD Mutations Relative to IL121_2014 |
| --- | --- |
| PDCoV RBD USA_IL121_2014 | N/A |
| PDCoV RBD HKU15_SD2018_300 | M354I, I391V, N397K |
| PDCoV RBD HKD_JPN_2016 | V326I, R342K |
| PDCoV RBD 110_1197_TW | V326I, T338S |
| PDCoV RBD Thai_S5011_2015 | M349L, T351R |

**Table S3.** CryoEM data collection and refinement statistics.

|  | PDCoV RBD with PD33 | PDCoV Spike SD2018/300 Apo | PDCoV Spike SD2018/300 with one PD41 bound | PDCoV Spike SD2018/300 with PD41 (local refinement) | PDCoV Spike SD2018/300 with two PD41 bound | PDCoV Spike SD2018/300 with three PD41 bound |
| --- | --- | --- | --- | --- | --- | --- |
| **Data collection and processing** |  |  |  |  |  |  |
| Magnification | 105,000 | 105,000 | 105,000 | 105,000 | 105,000 | 105,000 |
| Voltage (kV) | 300 | 300 | 300 | 300 | 300 | 300 |
| Electron exposure (e<sup>-</sup>/Å<sup>2</sup>) | 58 | 44.5 | 44.5 | 44.5 | 44.5 | 44.5 |
| Defocus range (μm) | 0.2 -3.0 | 0.8 -1.8 | 0.8 -1.8 | 0.8 -1.8 | 0.8 -1.8 | 0.8 -1.8 |
| Pixel size (Å) | 0.835 | 0.835 | 0.835 | 0.835 | 0.835 | 0.835 |
| Symmetry imposed | C1 | C3 | C1 | C1 | C1 | C1 |
| Final particle images (no.) | 564,616 | 24,642 | 465,175 | 465,175 | 99,272 | 26,452 |
| Map resolution (Å) | 3.0 | 3.1 | 2.72 | 3.0 | 3.0 | 3.4 |
| FSC threshold | 0.143 | 0.143 | 0.143 | 0.143 | 0.143 | 0.143 |
| Map sharpening B factor (Å<sup>2</sup>) | -139 | -55.3 | -85 | - 87.8 | -65.8 | -38.8 |
| **Validation** |  |  |  |  |  |  |
| MolProbity score | 1.73 |  |  | 0.87 |  |  |
| Clashscore | 3.17 |  |  | 0.2 |  |  |
| Poor rotamers (%) | 3.41 |  |  | 0 |  |  |
| Ramachandran plot |  |  |  |  |  |  |
| Favored (%) | 96.60 |  |  | 95.73 |  |  |
| Allowed (%) | 99.23 |  |  | 99.39 |  |  |
| Disallowed (%) | 0.77 |  |  | 0.61 |  |  |

## Resource availability

### Lead contact

Further information and requests for resources and reagents should be directed to and will be fulfilled by the lead contact, David Veesler.

### Materials availability

Materials generated in this study will be made available on request after signing a materials transfer agreement with the University of Washington. This work is licensed under a Creative CommonsAttribution 4.0 International (CC BY 4.0) license, which permits unrestricted use, distribution, and reproduction in any medium, provided the original work is properly cited. To view a copy of this license, visit https://creativecommons.org/licenses/by/4.0/. This license does not apply to figures/photos/artwork or other content included in the article that is credited to a third party;obtain authorization from the rights holder before using such material.

### Data and code availability

Data reported in this paper will be shared by the lead contact upon request. CryoEM maps and model data have been deposited at EMDB and PDB, respectively, and are publicly available as of the date of publication with accession numbers listed in the Key resources table. Raw sequencing data are available on the NCBI Sequence Read Archive and accession numbers are listed in the key resources table. DOIs are listed in the key resources table, figure legends, and method section. Any additional information required to reanalyze the data reported in this paper is available from the lead contact upon request.

## Experimental model and study participant details

### Cell lines

Cell lines used in this study consisted of DH10B competent cells (Thermo Fisher Scientific), BL21 DE3 (Sigma), HEK293T (ATCC, CRL-11268) and Expi293F (Thermo Fisher Scientific, A14527). HEK293T cells were cultured in 10% FBS (Fisher Scientific-Cytiva), 1% penicillin-streptomycin (Thermo Fisher Scientific) DMEM at 37°C, 5% CO2. Expi293F were cultured in expi293 media at 37°C, 8% CO2. None of the cell lines were authenticated or tested for mycoplasma contamination.

### *In vivo* animal studies

ATX-GK and ATX-GL female mice, 6-7 weeks old, were obtained from Alloy Therapeutics Inc. and housed for the immunization experiment at the Institute for Research in Biomedicine, Bellinzona, Switzerland. All animal experiments were performed in accordance with the Swiss Federal Veterinary Office guidelines and authorized by the Cantonal Veterinary (approval no. 35554 TI-39/2023/2023). Animals were supervised by a licensed veterinarian and proper steps were taken to ensure the welfare and minimize the suffering of all animals in the conducted studies. Animals were housed in ventilated cages in a 12 h light/dark cycle, with free access to water and standard sterilized chow.

## Method details

### PDCov immunization and antibody discovery from Alloy mice

Pre-immune serum was obtained from each mouse a week before immunization. ATX mice were successively immunized with recombinant PDCoV_IL121_2014_ RBD and S diluted (1:1) in Magic Mouse adjuvant (Cat#: CDN-A001E; CD Creative Diagnostics) and injected subcutaneously. On day 0, mice received prime immunization with 5 µg of PDCoV RBD_IL121_2014_ and were boosted on day 14 with 5 µg of PDCoV S_IL121_2014_ ectodomain. On day 29, the mice received 5 µg of PDCoV RBD_IL121_2014_ and on day 56 they received a last boost with 2 µg of PDCoV S_IL121_2014_. On day 64, the mice were sacrificed and peripheral blood, spleen and lymph nodes (LN) were collected and cells freshly isolated. B cells were enriched by positive selection using mouse CD19 microbeads and LS columns (Miltenyi). Enriched B cells were then stained with mouse anti-IgM, anti-IgD, and biotinylated PDCoV S_IL121_2014_ ectodomain labeled with both PE and Alexa-Fluor 647 streptavidin (Life Technologies). Sorted IgG+ memory B cells were seeded at clonal dilution in 384-well plates an a monolayer of feeder mesenchymal cells in the presence of B cell survival factors. Clones positive for antigen binding were then isolated, sequenced and produced recombinantly in transiently transfected CHO cells.

### Fab Fragment of monoclonal antibodies for binding and cryo-EM studies

Fab Fragments were generated by taking .5 mg of IgG and digesting with 2 µL of Lys C (NEB#P8109S) at 37℃. Digested product was bound to 500 µL of protein A resin beads (GenScript #L00210) overnight at 4℃ on a rotating shaker. Flow through post binding was collected, purified using size exclusion chromatography Superdex 200 10/300 and concentrated using Amicon Ultra-15 Centrifugal Filter Unit (10kDa). Purified Fab fragments were snap frozen and stored at -80℃.

### Production of anti-human kappa light chain nanobody

The pET23b with Anti-human kappa light chain nanobody with a C-terminal 8x His tag (PDB: 7WPD_Z) construct was transformed into *E. coli* BL21(DE3) cells. Overnight culture in Luria-Bertani (LB) media was initiated and incubated at 37 ℃ with shaking. 2L of LB media were inoculated with overnight culture and were induced in the exponential phase (OD600 = 0.6) by adding 1 mM IPTG. The cells were switched to 16 ℃ and grown overnight. Cells were collected by centrifugation for 15 minutes at 3000xg 4℃. Cell pellet was resuspended in 30 mL of cold 50mM Tris, 150mM NaCl, pH 8.0. Mixture was dounce homogenized 10x and sonicated using an Emulsiflex for 4x rounds. Cells were spun at 15,000 x g 4℃ for 30 minutes. Lysate was filtered using a 0.25 µm syringe filter and applied to Talon metal affinity resin (Takara) in 50mM Tris-HCl, 150mM NaCl, 10mM Imidazole at pH 8.0. Unbound proteins were washed in 10 column volumes of 50mM Tris-HCl, 150mM NaCl, 10mM Imidazole at pH 8.0. Eluted with 50mM Tris, 150mM NaCl, 250 mM Imidazole at pH 8.0. The eluted protein was concentrated using Amicon Ultra-15 Centrifugal Filter Unit (10kDa) and buffer exchanged into 50mM Tris, 150mM NaCl pH 8.0 and further purified using size exclusion chromatography Superdex 200 10/300. Nanobody proteins were snap frozen in liquid nitrogen and stored at -80℃.

### Production of recombinant PDCoV S glycoprotein

The PDCoV_IL121_2014_ S glycoprotein ectodomain (Genbank KJ481931.1) and SD2018/300 (Genbank KJ481931.1) were cloned into pcDNA3.1+ plasmid by GenScript with the host N-terminal signal peptide sequence, C-terminal foldon domain, thrombin cleavage sequence, short linker of GSG, AVI tag and 8 x His tag. The DNA constructs were expanded using DH5ɑ cells and purified using Qiagen MegaPrep Kit. Protein was expressed using ExpiFectamine 293 Transfection Kit (Thermo Fisher Scientific). Expi293F cells were grown at 37℃ with 8% CO_2_ to 3E6 cells and transfected with 100µg of DNA. Cell culture supernatants were harvested four days post-transfection. Protein was purified using HisTrap^Tm^ High Performance column (Cytiva). Protein was bound to HisTrap resin in 50mM Tris-HCl, 150mM NaCl, 10mM Imidazole at pH 8.0. Unbound proteins were washed in 10 column volumes of 50mM Tris-HCl, 150mM NaCl, 10mM Imidazole at pH 8.0. Eluted with 50mM Tris, 150mM NaCl, 250 mM Imidazole at pH 8.0. The eluted S protein was concentrated using Amicon Ultra-15 Centrifugal Filter Unit (100kDa) and buffer exchanged into 50mM Tris, 150mM NaCl pH 8.0. Endotoxin levels were assessed using Charles River Limulus Amebocyte Lysate (LAL) cartridges. Proteins were snap frozen in liquid nitrogen and stored at -80℃.

### Production of recombinant PDCoV RBD

The PDCoV_IL121_2014_ RBD (Genbank KJ481931.1) and SD2018/300 (Genbank KJ481931.1) spanning residues 303-415 were cloned into pcDNA3.1+ plasmid by GenScript with an N-terminal mu-phosphatase signal peptide sequence and C-terminal short linker GSG, AVI tag and 8 x His tag. The DNA constructs were expanded using DH5ɑ cells and purified using Qiagen MegaPrep Kit. Protein was expressed using ExpiFectamine 293 Transfection Kit (Thermo Fisher Scientific). Expi293F cells were grown at 37℃ with 8% CO_2_ to 3E6 cells and transfected with 100 µg of DNA. Cell culture supernatants were harvested four days post-transfection. Protein was purified using HisTrap^Tm^ High Performance column (Cytiva). Protein was bound to HisTrap resin in 50mM Tris-HCl, 150mM NaCl, 10mM Imidazole at pH 8.0. Unbound proteins were washed in 10 column volumes of 50mM Tris-HCl, 150mM NaCl, 10mM Imidazole at pH 8.0. Eluted with 50mM Tris-HCl, 150mM NaCl, 250 mM Imidazole at pH 8.0. The eluted RBD was concentrated using Amicon Ultra-15 Centrifugal Filter Unit (10 kDa) and buffer exchanged into 50mM Tris, 150mM NaCl pH 8.0 and further purified using size exclusion chromatography Superdex 200 10/300. Endotoxin levels were assessed using Charles River Limulus Amebocyte Lysate (LAL) cartridges. Proteins were snap frozen in liquid nitrogen and stored at -80℃.

### Production of recombinant APN ectodomain

The APN ectodomain from galline (Genbank ACZ95799.1) and human residues 66-967 was cloned into pcDNA3.1+ plasmid by GenScript with an N-terminal mu-phosphatase signal peptide sequence and C-terminal short linker GGS, thrombin cleavage site, human Fc fragment. The DNA constructs were expanded using DH5ɑ cells and purified using Qiagen MegaPrep Kit. Protein was expressed using ExpiFectamine 293 Transfection Kit (Thermo Fisher Scientific). Expi293F cells were grown at 37℃ with 8% CO_2_ to 3E6 cells and transfected with 100 µg of DNA. Cell culture supernatants were harvested four days post-transfection. Protein was purified using HighTrap Protein A column (Cytiva). Protein was bound to resin in 50mM Tris-HCl, 150mM NaCl pH 8.0. Unbound proteins were washed in 10 column volumes of 50mM Tris-HCl, 150mM NaCl at pH 8.0. Eluted with 0.1 M Citric Acid (pH 3.0) and neutralized with 1 M Tris-HCl (pH 9.0). The eluted APN was buffer exchanged into 50mM Tris, 150mM NaCl pH 8.0 and further purified using size exclusion chromatography Superdex 200 Increase 10/300 GL. Protein was concentrated using the Amicon Ultra-15 Centrifugal Filter Unit (100 kDa). Proteins were snap frozen in liquid nitrogen and stored at -80℃.

### Production of VSV pseudoviruses

PDCoV S glycoprotein constructs containing a C-terminal deletion of 21 residues to improve exportation to the membrane followed by a Flag tag were cloned into pcDNA3.1+ by GenScript. HEK293T cells were seeded at 16E6 cells in 100 mm dishes (Corning) poly-D-Lysine coated. Cells were transfected with 24 μg of DNA and 60 μL of Lipofectamine 2000 (Thermo Fisher Scientific) in Opti-MEM transfection medium. Post 5 hour incubation DMEM containing 20% FBS and 2% PenStrep was added to the cells. After 21 hours at 37 °C with 5% CO_2_ the cells are washed 1x with DMEM and infected with VSV (G∗ΔG-luciferase) in DMEM for 2 hours. The cells are washed 2x with DMEM and anti-VSV G antibody (I1-mouse hybridoma supernatant diluted 1:40 in DMEM, ATCC CRL-2700) is added and incubated for 24 hours at 37 °C with 5% CO_2_. Pseudovirus is harvested and collected by centrifugation at 3,000xg for 10 min. Supernatant containing the pseudovirus is filtered using a 0.45 μm syringe filter and concentrated 5x prior to storage at -80°C. Mock VSV pseudoviruses were prepared as above but without S glycoprotein transfection.

### Western Blot Analysis of VSV pseudoviruses

Western blot analysis was performed to determine the levels of S glycoprotein for each of the VSV pseudoviruses. 30 μL of virus is obtained and mixed with 10 μL of 4x LDS sample reducing buffer. Samples were boiled at 95 °C for 5 minutes and spun down. 35 μL of final mixture was run on 4-20% Mini-Protean TGX precast protein gel at 150 Volts for 45 minutes. Proteins were transferred using Trans-Blot Turbo Mini 0.2 µm PVDF Transfer Packs and membranes were incubated with Intercept (TBS) Blocking Buffer (Licor) for 1 hour at room temperature. Anti-Fusion peptide antibody at 1 μg/ml was diluted in Intercept (TBS) Antibody Diluent Buffer (Licor) and incubated at 4°C overnight. The following day membranes were washed with TBST 3x with 5-minute incubations at room temperature. Anti-human Fc (Jackson ImmunoResearch) was diluted 1:50,000 in Intercept (TBS) Antibody Diluent Buffer (Licor) and membranes were incubated at room temperature for 1 hour. Membranes were washed with 3x TBST with 5 minute incubations at room temperature. Images were taken using the Licor Odyssey CLx.

### Cell entry assays

HEK293T cells were transiently transfected with human APN (hAPN). hAPN with a C-terminal Flag tag was cloned into pcDNA3.1+ by GenScript. HEK293T cells seeded at 16E6 cells in 100 mm dishes coated with poly-D-Lysine and incubated overnight at 37 °C with 5% CO_2_. The following day cells were transfected with 8 μg of DNA using 30 μL of Lipofectamine 2000 transfection kit in Opti-MEM. Post 5 hours of incubation with transfection reagent, cells were trypsinized with 0.05% Trypsin EDTA for 3 minutes and neutralized with DMEM with 10% FBS 1% penicillin–streptomycin. 40,000 cells were seeded in 96-well plates (Corning 3610) coated with poly-lysine (Sigma P4707) and allowed to incubate at 37 °C with 5% CO_2_. Following day, media was removed and 20 μL of DMEM was added followed by 20 μL of VSV pseudotyped with PDCoV S variants for 2 hours at 37 °C with 5% CO_2_. Cells were supplemented with 40 μL of DMEM containing 20% FBS and 2% PenStrep and incubated for 22 hours at 37 °C with 5% CO_2_. 40 μL of One-Glo-EX substrate (Promega) was added and cells were incubated at 37 °C for 10 minutes with shaking. Luminescence was read using BioTek Neo2 plate reader. Technical replicates of three were run for three different lots of pseudovirus for each variant. Luciferase units were plotted using Graphpad Prism.

### Neutralization assays

For mAb neutralization assays against PDCoV S VSV, HEK293T cells were transfected with hAPN as described in the cell entry assay. 40,000 cells/well were seeded into 96-well plates coated with poly-lysine and incubated overnight at 37°C. Antibody titration series was performed in a half-area 96-well plate (Greiner) starting at a 2x concentration of 20 μg/ml and diluting 1:3 with DMEM. Equal volume of pseudovirus was added and incubated at room temperature for 30 minutes and 40 μL was transferred to cells and incubated for 2 hours at 37 °C with 5% CO_2_. Cells were supplemented with 40 μL of DMEM with 20% FBS and 2% PenStrep and incubated for 22 hours at 37 °C with 5% CO_2_. After 22 hrs, 40 μL of One-Glo-EX substrate (Promega) was added to each well and incubated on a plate shaker in the dark for 5 min before reading the relative luciferase units using a BioTek Neo2 plate reader. Relative luciferase units were used to determine (%) neutralization in Prism (GraphPad). Nonlinear regression curve fit and sigmoidal 4PL where x is the concentration was run to determine IC50 values for 3 biological replicates of pseudovirus and three technical replicates for each titration series.

### Biolayer interferometry analysis (BLI)

BLI binding assays were performed on an Octet Red (Sartorius) instrument operated at 30°C with shaking (1000 rpm). Nickel-NTA biosensor tips were hydrated in 10 x Kinetics Buffer for 10 minutes at room temperature. PDCoV RBD-His was loaded at 10 μg/ml in 10 x Kinetics Buffer to 1 nm. Loaded tips were dipped into a 10x Kinetics buffer to stabilize and remove any unbound protein for 60 seconds. Loaded RBD tips were dipped into 1 μg/ml of Fab fragment for 360 seconds. Associated Fab fragment to the RBD was preceded by a dissociation phase of tips being moved into a 10x kinetics buffer for 360 seconds. K_D_ determination of PD33 and PD41 Fab to RBD was obtained through a concentration series starting at 25 nM and titrated down by two fold down to 0.39 nM. KD_apparent_ determination of RBD to hAPN-Fc was obtained through a concentration series starting at 5 μM and titrated down by two fold down to 39 nM. K_D_ values were calculated using ForteBio data analysis software where fitting of curves using a 1:1 binding model was used. For the competition experiment of Fab and gAPN-Fc post loading with RBD, 25nM of Fab was tested against 250nM of gAPN-Fc. For the competition experiment of Fab PD33 and Fab PD41, PDCoV RBD-AVI-Biotinylated was loaded at 2.5 μg/ml in 10 x Kinetics Buffer to 1 nm, 1 μg/ml of associated and bound Fab PD33 was tested against 1.0 μg/ml of Fab PD41 and the opposite was performed where 1 μg/ml of Fab PD41 was tested against 1.0 μg/ml of Fab PD33.

### Cryo-EM sample preparation and data collection

For PD33 Fab bound PDCoV RBD structure, 2 μM PDCoV RBD complex with 1 μM PD33 Fab and 1.5 μM anti-human kappa light chain nanobody were prepared by incubation for 1 hour at 4°C. Nanobody was used to stabilize the constant region of the PD33 Fab structure. 3ul of PDCoV RBD/PD33Fab/nanobody complex were directly applied to freshly glow-discharged grids. The Cryo-EM grids were prepared using a Vitrobot Mark IV (Thermo Fisher Scientific) with R 2/2 UltrAuFoil grids and a Chameleon^55^ (SPT Labtech) with self-wicking nanowire Cu R1.2/0.8 holey carbon grids. Data were acquired using an FEI Titan Krios transmission electron microscope operated at 300 kV and equipped with a Gatan K3 direct detector and Gatan Quantum GIF energy filter operated in zero-loss mode with a slit width of 20 eV. Automated data collection was carried out using Leginon^56^ at a nominal magnification of 105,000 with a pixel size of 0.835 Å and total exposure dose of 58 e/A^2^. 13,767 micrographs from UltrAuFoil^57^ grids and 13,733 micrographs from chameleon grids were collected with a defocus range between -0.2 and -3.0 μm. Each movie was fractionated in 100 frames of 40 ms each.

Cryo-EM grids of 1.5 μM PDCoV S SD2018/300 complexed with 4.5 μM of PD41 Fab for 1 hour at 4°C were added to glow discharged 300 1.2/1.3 grids. The grids were blotted with a blot force of -1, 6 second blot time and 10 second wait time before plunge freezing using a vitrobot MarkIV (ThermoFisher Scientific) set at 100% humidity and 4 °C. Data were acquired using an FEI Titan Krios transmission electron microscope operated at 300 kV and equipped with a Gatan K3 direct detector and Gatan Quantum GIF energy filter operated in zero-loss mode with a slit width of 20 eV. Automated data collection was carried out using SerialEM^58^ at a nominal magnification of 105,000 with a pixel size of 0.835 Å in defocus range between -1.8 to -0.8 μm. Total exposure dose of 44.5 e/A2. Each movie was fractionated in 79 frames of 40 ms each. A total of 19,556 micrographs were collected.

### Cryo-EM data processing, model building and refinement

The raw movie data of PDCoV RBD complexed with PD33 Fab and anti-human kappa light chain nanobody were processed using motion correction, contrast-transfer function (CTF) parameter estimation, automatic particle picking, and extraction using Warp^45^. Particles were extracted with a box size of 168 pixels with a pixel size of 1.67Å. After two rounds of 2D classification using cryoSPARC^46^, well-defined particles were selected and particles from each dataset were combined. Initial models were generated with ab-initio reconstruction in cryoSPARC. The initial models were used as references for a heterogenous refinement. The Topaz models were trained on a Warp particle set which did not yield a high-resolution reconstruction. The particles picked using Topaz were extracted and particles were subjected to 2D-classification, ab-initio reconstruction and heterogenous 3D refinement in cryoSPARC. Particle picking with Topaz improved the number of unique 2D views. The two different particle sets from the Warp and Topaz strategies were merged and duplicates were removed. After two rounds of 3D heterogeneous refinements and removal of junk particles, 3D refinements were carried out using non-uniform refinement. Particle data sets transfer between RELION and cryoSPARC was performed by the pyem program package^47^. Selected particle images were subjected to the Bayesian polishing procedure implemented in RELION^48^. To further improve the density of the PD33 Fab and RBD, the resulting particle sets were subjected to cryoSPARC 2D-classification, ab-initio reconstruction and 3D heterogeneous refinement. Particles belonging to classes with the best resolved PD33 Fab and RBD density were selected and subjected to final 3D non-uniform refinement using cryoSPARC to yield the final reconstruction at 2.9 Å resolution constituting 564,616 particles.

PDCoV SD2018/300 with PD41 Fab data was processed using RELION motion correction, contrast-transfer function (CTF) parameter estimation, particle picking, and extraction using cryoSPARC. Particle images were extracted with a box size of 512 downsampled to 256 pixels and a pixel size of 0.828Å (longest axis for S with Fab is 206 Å). After two rounds of 2D classification using cryoSPARC, well-defined 465,175 particles were selected and processed through non-uniform refinement (NUR). Particles were separated into different 3D classes using 3D classification in RELION with 50 iterations (angular sampling of 7.5 angular sampling 7.5 °; for 25 iterations and 1.8°; with local search for 25 iterations). Particles were grouped into four 3D classes; Class I) apo II) one RBD bound to PD41 III) two RBDs bound to PD41 and IV) three RBDs bound to PD41. Class I, II, IV particles individually were imported into cryoSPARC and refined using NUR in cryoSPARC. Imported particles from class I, II, III, IV were combined for solving a higher resolution structure of the RBD:PD41 and subjected to NUR in cryoSPARC. Local refinement was performed with a mask around the RBD: PD41 region generated from a volume with a threshold of 0.01 and soft padding width of 17 pixels.

Reported resolutions are based on the gold-standard Fourier shell correlation (FSC) of 0.143 criterion. Local resolution estimation, filtering, and sharpening were carried out using cryoSPARC. The resulting map showed clear density for the overall quaternary architecture and side chains of the interface of PDCoV RBD and PD33 Fab complex. Chimera, ChimeraX and Coot were used to fit atomic models into the cryo-EM maps^49–51^. Structure was refined and relaxed using Rosetta using sharpened and unsharpened maps and validated using Phenix and Molprobity^52–53^. Analysis of interface residues was assisted by PISA.

### Quantification and statistical analysis

Quantification details of experiments can be found in the figure legends.

## Acknowledgements

This study was supported by the National Institute of Allergy and Infectious Diseases (P01AI167966 to T.N.S., and D.V., DP1AI158186 and 75N93022C00036 to D.V.), a Pew Biomedical Scholars Award (D.V.), an Investigators in the Pathogenesis of Infectious Disease Awards from the Burroughs Wellcome Fund (D.V.), a Dale F. Frey Award for Breakthrough Scientists from the Damon Runyon Cancer Research Foundation (T.N.S.), the University of Washington Arnold and Mabel Beckman Cryo-EM Center, the National Institute of Health grant S10OD032290 (to D.V.), the National Science Foundation Graduate Research Fellowship (NSF-GRFP DGE-2140004 to M.R.). D.V. is an investigator of the Howard Hughes Medical Institute and the Hans Neurath Endowed Chair in Biochemistry at the University of Washington.

## Author contributions

M.R., L.P., F.B., and D.V. designed the experiments. M.R., L.P. and J.T.B. recombinantly expressed and purified glycoproteins. M.R. purified and characterized the Fabs and anti-kappa light chain nanobody. L.P., A.D., and F.B. performed immunization and monoclonal antibody isolations. K.C., C.S. and A.B. produced the recombinant antibodies. M.R. performed binding assays, produced pseudovirus and performed entry assays. K.S. and C.N.Y. helped with generating hybridoma and parent pseudovirus. Y.J.P. carried out cryo-EM specimen preparation, data collection, and processing of the PD33-bound PDCoV RBD structure. M.R. carried out cryo-EM specimen preparation, data collection, and processing of the PD41-bound PDCoV S_SD2018/300_. Y.J.P., D.A., S.D., J.Q., and D.V. helped with cryoEM characterization of the PD41-bound PDCoV S_SD2018/300_. M.R., Y.J.P., and D.V. build and refine atomic models. M.M. cloned the construct and produced an initial batch of the anti-kappa light chain nanobody. A.T. provided the constructs for human APN. A.T. and T.S. performed bioinformatic analysis to aid in strain selection of PDCoV. M.R., Y.J.P. and D.V. analyzed the data and wrote the manuscript with input from all authors. D.C., F.B. and D.V. supervised the project.

## Declaration of interests

D.V. is named as inventors on patents for coronavirus nanoparticle vaccines filed by the University of Washington. D.C., L.P., A.D., K.C., C.S., A.B., and F.B. are employees of Vir Biotechnology and may hold shares in Vir Biotechnology. The remaining authors declare that the research was conducted in the absence of any commercial or financial relationships that could be construed as a potential conflict of interest.

## Notes

### Summary of Updates

Comments were visible from the Word document

